# Small silencing RNAs expressed from W-linked retrocopies of *Masculinizer* target the male-determining gene *PxyMasc* during female sex determination in the Diamondback moth *Plutella xylostella*

**DOI:** 10.1101/2022.04.04.486979

**Authors:** T. Harvey-Samuel, X. Xu, M. A. E. Anderson, L. Carabajal Paladino, D. Kumar Purusothaman, V.C. Norman, C.M. Reitmayer, M. You, L. Alphey

## Abstract

The Lepidoptera are an insect order of cultural, economic and environmental importance, representing c. 10% of all described living species. Yet, for all but one of these species (silkmoth, *Bombyx mori*) the molecular genetics of how sexual fate is determined remains unknown. We investigated this in the diamondback moth (DBM - *Plutella xylostella*), a globally important, highly invasive and economically damaging pest of cruciferous crops. Our previous work uncovered a regulator of male sex determination in DBM – *PxyMasc*, a homologue of *B. mori Masculinizer* - which although initially expressed in embryos of both sexes, is then reduced in female embryos, leading to female-specific splicing of *doublesex*. Here, through sequencing small RNA libraries generated from early embryos and sexed larval pools, we identified a variety of small silencing RNAs (predominantly piRNAs) complementary to *PxyMasc*, whose temporal expression correlated with the reduction in *PxyMasc* transcript observed previously in females. Analysis of these small RNAs showed that they are expressed from tandemly-arranged, multi-copy arrays found exclusively on the W (female-specific) chromosome, which we term ‘*Pxyfem*’. Analysis of the *Pxyfem* sequences showed that they are partial cDNAs of *PxyMasc* mRNA transcripts, likely integrated into transposable element graveyards by the non-canonical action of retrotransposons (retrocopies), and that their apparent similarity to *B. mori feminizer* more probably represents convergent evolution. Our study helps elucidate the sex determination cascade in this globally important pest and highlights the ‘shortcuts’ which retrotransposition events can facilitate in the evolution of complex molecular cascades, including sex determination.

**Significance statement:** Uncovering the mechanisms which species have evolved to determine sex is of fundamental interest and provides avenues for pest management through genetic manipulation of these pathways. In insects, much of what is known regarding sex determination is concentrated within the Diptera and Hymenoptera, despite other orders (e.g. Lepidoptera) being of great ecological and economic importance. Here, using small RNA sequencing of embryonic and early larval samples, we uncover an RNAi-based sex determination system which silences the male determining gene *PxyMasc* in the Diamondback moth (*Plutella xylostella*) – a global pest of cruciferous crops. We track production of these silencing RNAs back to the W-chromosome where they are expressed from partial cDNA copies of *PxyMasc*. Our analysis suggests these are *PxyMasc* ‘retrocopies’, integrated via the non-canonical action of LTR retrotransposons and that similarities between this system and the *feminizer* system in *Bombyx mori* likely represent convergent evolution.

## Introduction

Insects have evolved a diverse range of mechanisms to determine sex, which vary widely both between and within orders (1). Even within groups where the broad pattern of sex determination is similar (e.g. XY males / XX females), this can be achieved through non-homologous molecular mechanisms, for example X-linked gene dosage or dominant male determining loci (2). The study of these mechanisms is of fundamental interest, but also provides tools and targets for Genetic Pest Management (GPM) strategies, including gene drives (3, 4).

Much of what is known regarding insect sex determination systems is based on examples from a few orders of economic or human health importance, primarily the Diptera and Hymenoptera (2). The Lepidoptera, which contain approximately 10% of all described living species, including many of economic, cultural and environmental importance (5), are understudied in this regard – only the domestic silkworm (*Bombyx mori*) has had the molecular mechanism by which it determines sex uncovered (6). In common with other lepidopterans, *B. mori* exhibits female heterogamety (WZ/ZZ females/males, although Z0/ZZ systems are present in basal Lepidoptera) (7). Recent work identified the master regulator of sex in *B. mori* to be a single piRNA (known as *feminizer - fem*) complementary to the Z-linked male determining gene *Masculinizer* (6). This 29bp ‘*fem*’ piRNA exists in multiple copies, exclusively on the female-specific W-chromosome. In the presence of the W-chromosome (i.e. in females), *Masculinizer* mRNA is silenced by *fem*, allowing the female sex-determination cascade to engage. In the absence of the W-chromosome (i.e. in ZZ males), *Masculinizer* is translated, initiating the male cascade (6).

As the first recorded example of a RNAi-controlled sex determination cascade and the first uncovered cascade in any lepidopteran, this study was of great significance. However, questions regarding its generality remain. For example, the W-chromosome is known to have evolved independently at least twice within the Lepidoptera (Z0/ZZ being the ancestral arrangement) with multiple examples of neo-Z fusions in the upper Ditrysian Lepidoptera (8) suggesting that the true number of independent W-generating events may be higher (9, 10). It thus remains unclear to what extent the W-linked *feminizer* system uncovered in *B. mori* is conserved, especially given this species’ long history of domestication (c. 5,000 years of artificial rearing) and the fact that at least one wild silkmoth species (*Samia cynthia*), although possessing a W-chromosome, does not require it for female sex determination (11). Furthermore, although the function and Z-linkage of *Masculinizer* appear to be deeply conserved, examples identified so far show that the amino acid sequence of this gene has diverged significantly over evolutionary time (12), to the extent that it is unclear how a multi-copy, nucleotide-specific silencing RNA would have been able to maintain sequence complementarity. Indeed the nucleotide sequence targeted by *fem* is not conserved even in the closely related *Trilocha varians Masculinizer* homologue (13). Finally, to our knowledge, no hypotheses have been proposed for how such a unique mechanism of sex-determination may have initially evolved.

We chose to explore these questions in the diamondback moth (DBM) *Plutella xylostella*. In addition to being one of the world’s most damaging agricultural pests (14-16) and one in which GPM strategies are being developed (17-20), DBM possesses characteristics which are beneficial in addressing these questions. It is a member of the superfamily Yponomeutoidea, an early-diverging Ditrysian lineage evolutionarily distant from the more derived Bombycoidea (which includes *B. mori*) and thus important in exploring early lepidopteran evolution (9). It possesses the ancestral lepidopteran chromosome number (n=31), with no large-scale chromosome rearrangements and a well-differentiated WZ/ZZ sex-chromosome system (9, 21, 22). Finally, it has been subjected to an unusually high degree of molecular genetic research for a lepidopteran, with publicly available male- and female-derived draft genome sequences (21, 23), RAD-based chromosome-level maps (24), extensive karyotyping analysis and, crucially, a recently identified and characterised *Masculinizer* homologue (*PxyMasc*) (25).

Here, using the *PxyMasc* mRNA sequence as a guide, we identified a functionally conserved yet likely independently derived *fem*-like system in DBM. We observed a range of small RNAs, expressed from early embryonic stages onwards, mapping to both the sense and antisense strands of *PxyMasc* exons four, five and six, including size ranges corresponding to both siRNA/miRNAs and piRNAs. Rapid amplification of cDNA ends (RACE) conducted against these antisense piRNAs identified transcripts consisting of contiguous reverse complement *PxyMasc* and truncated retrotransposon ORF sequences. These putative *P. xylostella fem* (‘*Pxyfem*’) precursors mapped to multi-copy loci within ‘transposable element graveyards’ found exclusively in a female genome assembly and which genomic PCR confirmed were female-specific (i.e. W-linked). Intriguingly, analysis of the genomic context of these sequences suggests that *Pxyfem* evolved through a template switching event involving Long Terminal Repeat (LTR) retrotransposons and a *PxyMasc* mRNA transcript, followed by retroposition of the chimeric RNA onto the W-chromosome. Analysis of the ‘fossilised’ W-linked LTR remnants and the *Pxyfem* sequences themselves suggests that this event took place after the divergence of *P. xylostella* from other Ditrysians whose genomes have been sequenced, suggesting independent evolution of this female *Masculinizer* regulator relative to that of *B. mori*.

## Results & Discussion

### A range of small RNAs map to *PxyMasc*

If a system similar to *B. mori fem* existed in DBM, we would expect small RNA libraries (reads of 18-40 nucleotides in length) to produce sequences mapping to the antisense strand of *PxyMasc* in female-but not male-derived pools. Out of 3,328,300 female L1 larvae derived small RNA reads, 38 were found to map to the *PxyMasc* mRNA sequence, with 12 of these mapping to the sense and 26 to the antisense strand. For the male L1 larvae derived RNA, of 2,369,543 reads, only two mapped to *PxyMasc* with neither of these mapping to the antisense strand (Figure 1).

**Figure 1.**
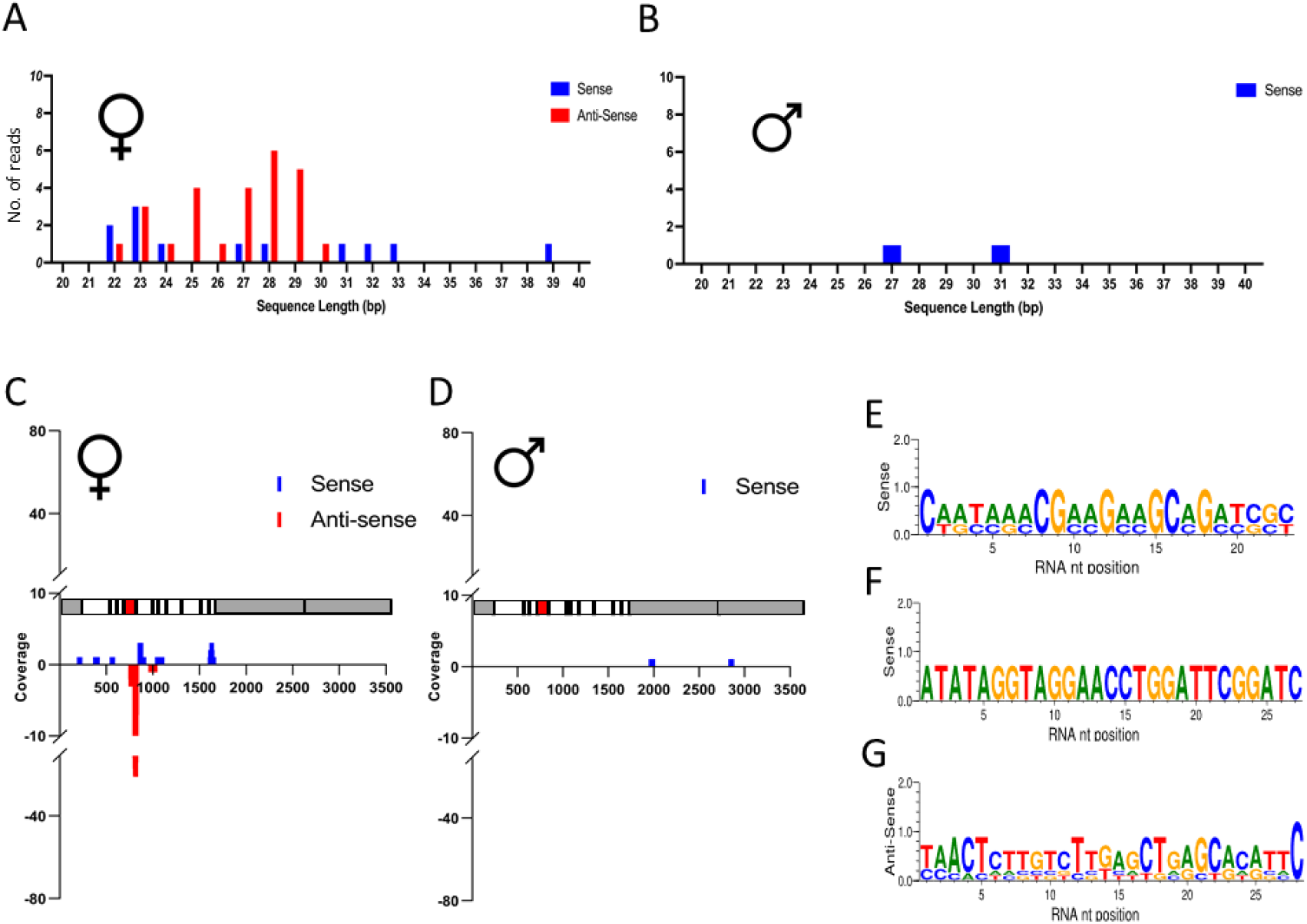
Small RNA deep sequencing of female or male pooled L1 samples: A, C, E, G: Data deriving from L1 female pools. B, D, F: Data deriving from L1 male pools. A and B: number, size and orientation of reads mapping to the *PxyMasc* mRNA sequence. C and D: location along *PxyMasc* mRNA sequence of mapped reads. A schematic of the *PxyMasc* mRNA is shown parallel with each X axis to show features against which these reads map. Grey areas designate UTRs, white areas designate coding exons, the red shaded area designates the region of the transcript coding for the Cysteine-Cysteine motif – a domain functionally requisite in *Bombyx mori* Masc and conserved amongst other lepidopteran Masc homologues identified to date. E, F, G: sequence logos for the most common length of sense and antisense reads identified in the two pools. Note: the male pool did not provide any antisense reads mapping to *PxyMasc*.

For the female L1 derived library, the majority of reads mapped to *PxyMasc* exons 5 and 6, with a particular concentration of reads mapping to the antisense strand of the exon 5-6 junction (Figure 1C), overlapping with the highly conserved Cysteine-Cysteine domain - a hallmark of identified Masculinizer homologues and an area required for functionality in *B. mori* Masc (26). The two male reads mapped to independent sequences within the putative 3’UTR of *PxyMasc*. These two reads were not present in the female-derived library (Figure 1D).

Interestingly, unlike the *B. mori fem* transcript which produces a single, 29bp piRNA, in the female L1 derived sample we observed multiple different *PxyMasc* mapping reads whose size ranges corresponded to both siRNA/miRNAs (∼21-24bp) and piRNAs (∼25-30bp), although those belonging to the ‘piRNA’ size class predominated (Figure 1A). We did not observe the classic ‘ping-pong’ signal associated with these reads (3’ overlap of 10bp between forward and reverse reads), however, they did display a bias towards a 5’ end uridine (1U-bias) in antisense strand mapping reads, a signal of piRNA biogenesis (27) (Figure 1G).

Subsequently, we set out to establish whether these *PxyMasc*-mapping reads were present in early embryos, as would be expected of a *fem*-like sex determination system. Previously, we established that sex in DBM is determined in the early embryo (6-24h after laying) by assessing the temporal expression pattern of *PxyMasc* and *doublesex* during early embryos (25). There, *PxyMasc* mRNA was not observed at early time points (3h post oviposition). However, by 6h post oviposition, *PxyMasc* mRNA was present in both sexes and being translated, judging by observed male-form splicing of *doublesex* (a function of PxyMasc, or its downstream cascade). By 24h post oviposition, *PxyMasc* mRNA was no longer detectable in female embryos, but remained present in males. This implied that the silencing of *PxyMasc* by any putative *fem*-like system would likely begin and peak during the 6-24h period post oviposition. To assess this, we sequenced pooled embryonic small RNA libraries at 3, 6, 9, 12 and 24h post oviposition (Figure 2). As expected, the 3h library did not contain *PxyMasc* mapping reads. However, from 6h onwards, such reads were observed, gradually increasing to the 12h time point, before dropping away by 24h (Supplementary Table 1 for total and normalised read numbers). The position of these reads generally concurred with those observed in the female L1 derived sample and, again, read lengths corresponding to siRNA/miRNAs as well as piRNAs were observed. As with the female L1 sample, we observed a 1U-bias in antisense mapping reads only. However, we now also observed a 10A-bias, in sense mapping reads only (see Figure 2 for demonstration of this in the most common size class of read in each timepoint). Additionally, sense and antisense reads showing a 3’ 10bp overlap ‘ping-pong’ signal (28) were apparent - see Figure 3 for closer examination of this in the 12h sample. Taken together, these data indicate a ping-pong amplification cycle of piRNAs targeting *PxyMasc* early in DBM embryonic development. Interestingly, secondary concentrations of sense and antisense reads mapped down and upstream, respectively, of the primary exon 5-6 junction concentration (see Figure 3), which may be indicative of a ‘phased’ piRNA biogenesis (29, 30) - where the primary transcript these piRNAs derived from is serially processed into downstream piRNAs after the initial cleavage event. Further interrogation of the *Zucchini*-dependent processing pathway would be required to verify this.

**Figure 2:**
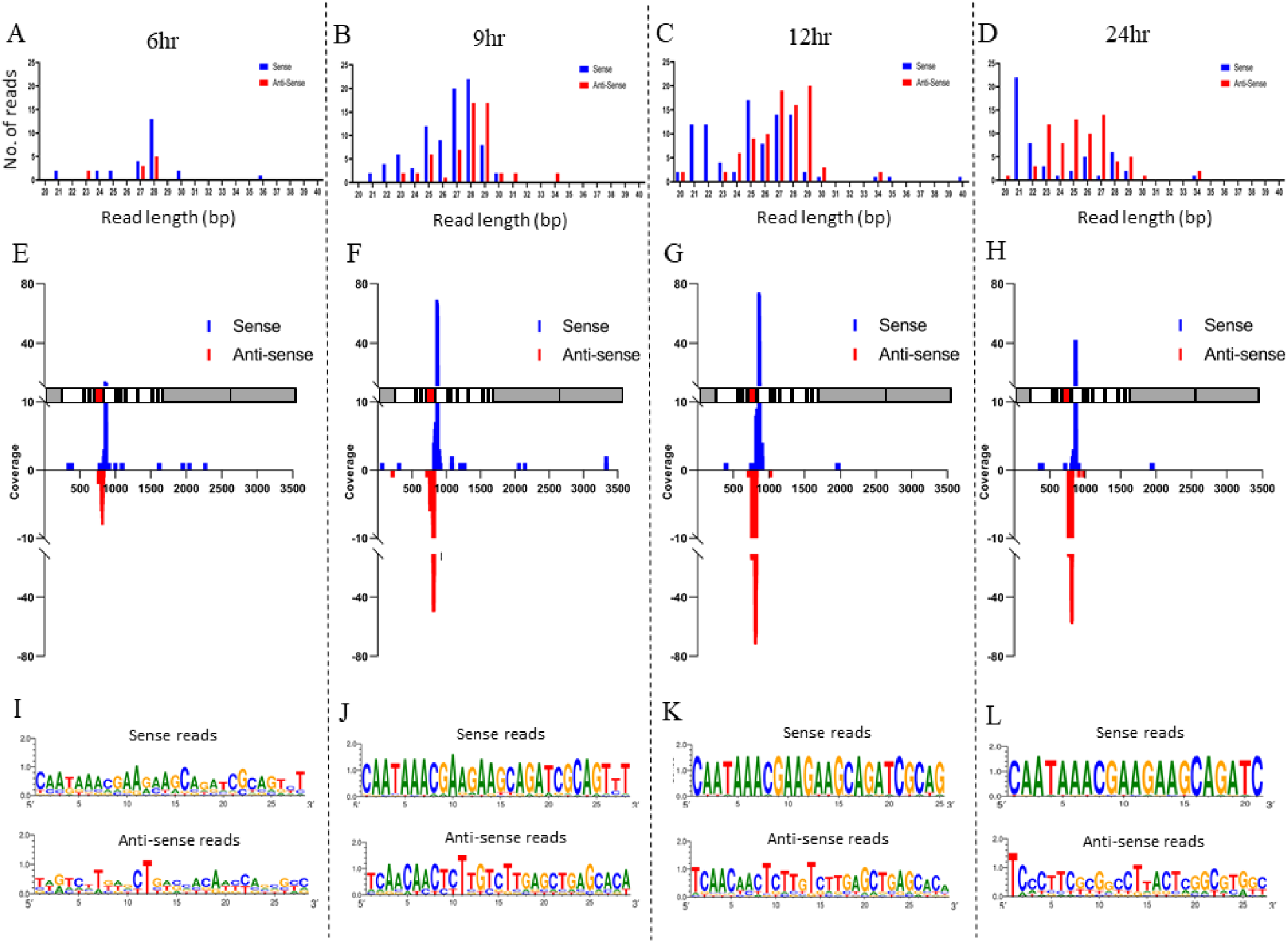
Small RNA deep sequencing of 6, 9, 12, and 24h pooled embryo samples: A, E, I: data deriving from 6h pooled embryos. B, F, J: data deriving from 9h pooled embryos. C, G, K: data deriving from 12h pooled embryos. D, H, L: data deriving from 24h pooled embryos. A, B, C, D: number, size and orientation of reads mapping to the *PxyMasc* mRNA sequence. E, F, G, H: location along *PxyMasc* mRNA sequence of mapped reads. A schematic of the *PxyMasc* mRNA is shown parallel with each X axis to show features against which these reads map. Grey areas designate UTRs, white areas designate coding exons, the red shaded area designates the region of the transcript coding for the Cysteine-Cysteine motif – a domain functionally requisite in *Bombyx mori* Masc and conserved amongst other lepidopteran Masc homologues identified to date. I, J, K, L: sequence logos for the most common length of sense and antisense reads identified in the two pools.

**Figure 3:**
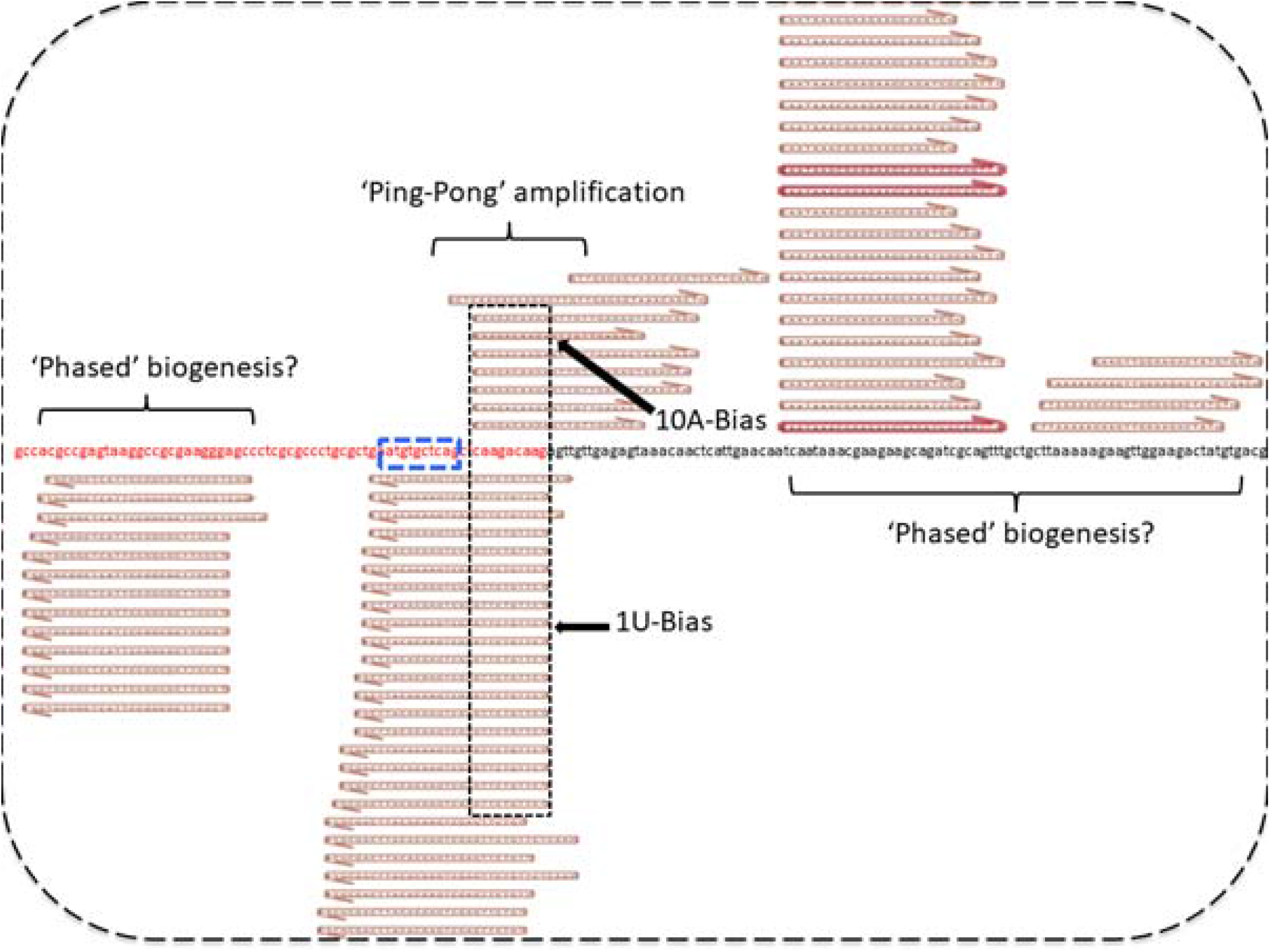
Ping-pong amplification signal evident in pooled 12h embryo *PxyMasc* mapping reads. Central nucleotide sequence is of the *PxyMasc* mRNA exon 5 (shown in red lettering) and exon 6 (shown in black lettering) junction. Blue dashed box designates the nucleotide sequence translated into the Cysteine-Cysteine domain. Sense and antisense reads mapping to this area show characteristic signs of the ‘ping-pong’ amplification loop, overlapping on their 3’ ends by 10bp with a 1-U bias for antisense reads and 10-A bias for sense reads. Reads mapping away from this central area showed a strong bias for mapping in the same direction as those in the ping pong cycle e.g. reads shown mapping in exon 5 were all in an antisense orientation. This may be indicative of a ‘phased’ biogenesis for these small silencing RNAs.

These trends concur with the observed patterns of *PxyMasc* and sex-specific *doublesex* mRNA presence in males and females described previously (25)and suggest a potential role for these small silencing RNAs (ssRNAs) in mediating that pattern.

### Antisense ssRNAs derive from transcripts associated with the W-chromosome

To identify the primary transcripts from which these *PxyMasc*-targeting ssRNAs were derived, 5’ and 3’ RACE gene specific primers (GSPs) were designed in a region of exon 5 where the largest concentration of those reads were observed. Using GSPs matching the sense sequence of *PxyMasc* for 5’ RACE (and the antisense sequence of *PxyMasc* for 3’ RACE) prevented the unintentional amplification of the endogenous *PxyMasc* transcript. One 5’ and one 3’ transcript were amplified from ovarian RACE-ready cDNA. The 5’ RACE transcript consisted of a cDNA copy of *PxyMasc* extending upstream from the primer binding point (in exon 5) into exon 6 (i.e. a partial reverse complement - RC – of *PxyMasc* spliced mRNA). BLASTn analysis of the remainder of the transcript revealed a 490bp sequence (henceforth Multiply Repeated Region 1 - MRR1) showing high similarity (highest=97%) to multiple (>80) hits in both the male-derived DBM genome assembly (PRJNA277936) and female-derived DBM genome assembly (PRJEB34571) (Figure 4A). However, the full RACE transcript sequence was found only in the female-derived assembly. Similarly, the 3’ RACE product consisted of RC-*PxyMasc* exons 5 and 4 (Figure 4B). The full-length RACE transcript sequence could again only be found in the female-derived assembly, whereas a shorter 513bp region of the transcript showed high similarity (99.4%) to multiple repeated regions in both the male and female-derived genome assemblies (henceforth Multiply Repeated Region 2 - MRR2).

**Figure 4:**
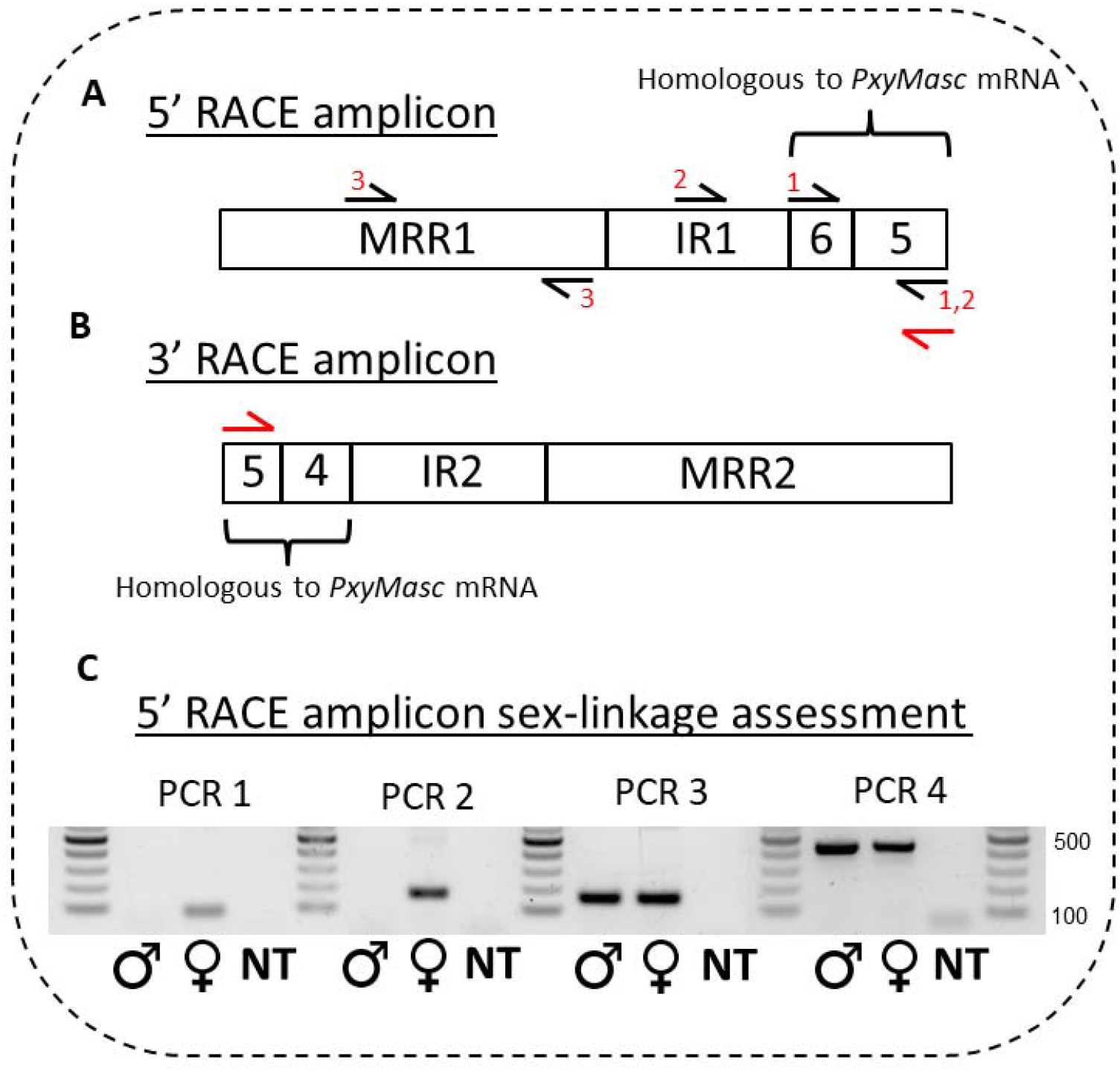
Schematic of observed RACE transcripts and PCR assessment of RACE amplicon sex linkage. A: Schematic of 5’ RACE amplicon observed when using a gene specific primer in the sense orientation to *PxyMasc* exon 5 (shown by red primer symbol). MRR1 = Multiply Repeated Region 1, IR1 = Intervening Region 1, 5 and 6 = sequence homologous to *PxyMasc* exons 5 and 6. Black primer symbols and respective numbers listed show the approximate locations and combinations of primers used in PCRs listed in (C). B: Schematic of 3’ RACE amplicon observed when using a gene specific primer in the antisense orientation to *PxyMasc* exon 5 (shown by red primer symbol). MRR2 = Multiply Repeated Region 2, IR2 = Intervening Region 2, 4 and 5 = sequence homologous to *PxyMasc* exons 4 and 5. C: PCR assessment of sex-linkage of the genomic locus from which the 5’ RACE amplicon was expressed. The same number above wells denotes a consistent primer set used in that PCR. Position of primer sets 1,2 and 3 shown in A. + control (PCR 4) shows amplification of the 17S gene.

We hypothesised that in both cases the full RACE transcripts were identified only in the female-derived assembly as they were located on (expressed from) the W-chromosome, whereas the MRRs may represent sections of transposable elements present on the W-chromosome but also elsewhere in autosomal regions (and therefore also in the male-derived assembly). To test this we performed a series of genomic PCRs across the 5’ RACE transcript sequence (Figure 4A). PCR amplicons were produced within the MRR1 region when both male and female gDNA samples were used as templates. However, amplicons across the MRR1/*PxyMasc* fragment junction were only produced when female gDNA was used, implying that these regions are exclusive to the female genome and therefore W-linked (Figure 4C).

### *Pxyfem* occurs in two multi-copy tandemly arranged clusters within transposable element graveyards

BLASTn analysis showed that sequences highly similar to the full-length RACE transcripts mapped exclusively to a female-derived genomic scaffold (CABWKK010000004) with no matches in the male-derived genome assembly. Analysis of this scaffold uncovered 11 loci with sequences similar to the *PxyMasc* CDS (henceforth ‘*Pxyfem*’ loci), in two divergently orientated clusters (Figure 5A). Whilst all 11 *Pxyfem* loci included *PxyMasc* exons (partial) 4, 5 and 6, 7 of these continued partway into exon 7 (henceforth ‘<u>long</u>’ *Pxyfem*=326bp) whilst 4 ended partway through exon 6 (henceforth ‘<u>short</u>’ *Pxyfem*=192bp) (Figure 5B).These *Pxyfem* loci showed very high nucleotide sequence conservation both to *PxyMasc* (average similarity to *PxyMasc*=93.46% +/- 0.26) and to each other (92.7% sequence conservation between all loci), calculated over the core 192bp region shared by both short and long *Pxyfem*. RT-PCR using ovarian-derived cDNA and primers which orientated ‘outwards’ from each *Pxyfem* locus showed that at least some closely linked *Pxyfem* loci are expressed on the same transcript (Supplementary Figure 2).

**Figure 5:**
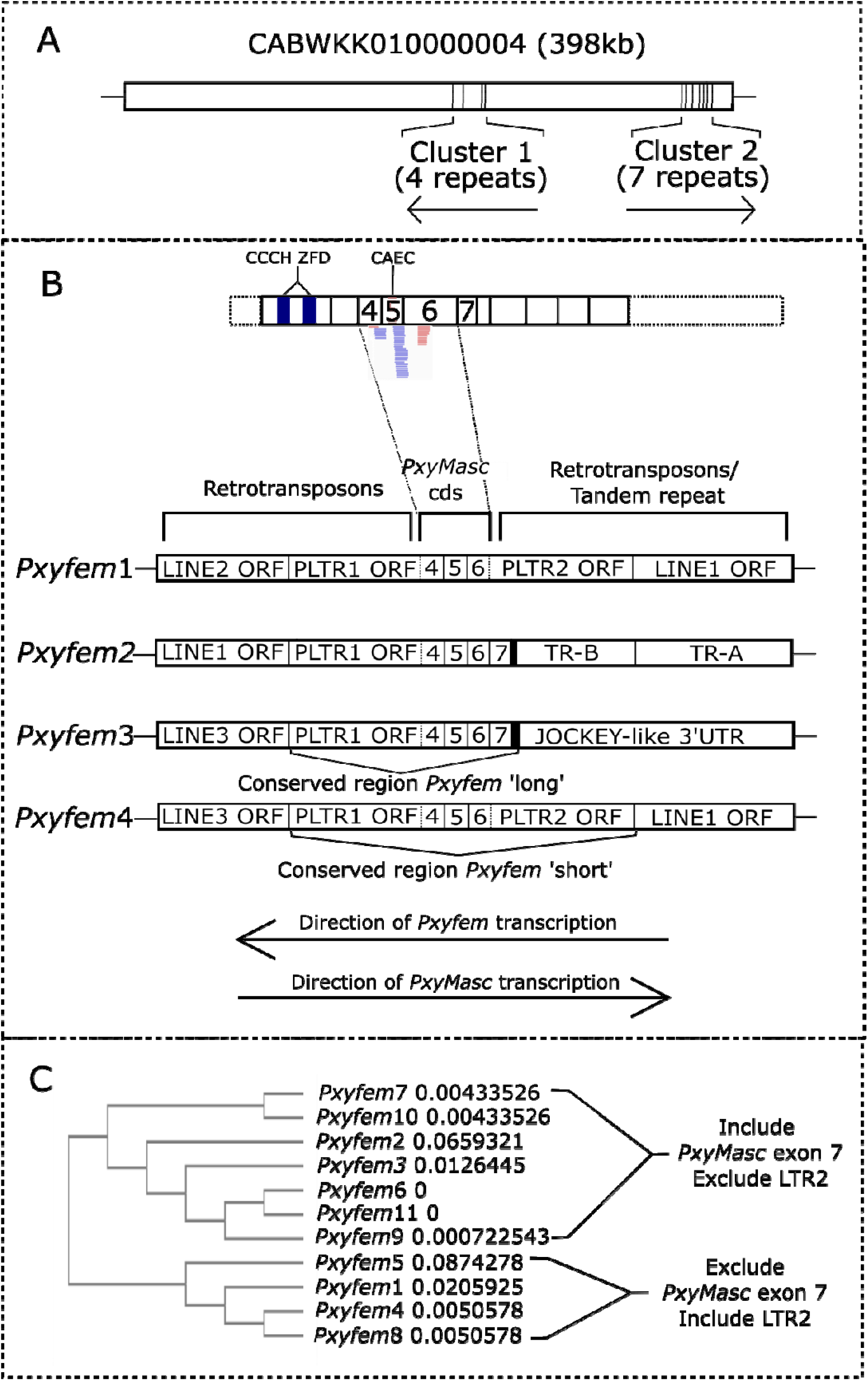
RACE transcripts are derived from genomic loci nested within transposable element graveyards. A: Full length 5’ and 3’ RACE amplicon sequences mapped exclusively to a scaffold from the female-derived DBM genome assembly (scaffold CABWKK010000004 from genome project PRJEB34571). These ‘*Pxyfem*’ loci/copies occurred in two, differentially orientated clusters within each of which, copies were tandemly arranged. Orientation of the copies shown by arrows under each cluster. B: Schematic showing detail of the *Pxyfem* loci included in Cluster 1 as well as c. 500bp of up and downstream genomic flanking sequence. Identity of truncated genes within these flanking sequences were elucidated through ORF analysis and BLASTp, or BLASTn of the nucleotide sequence if no homologous ORFs were observed. Schematic above these four copies is of the full *PxyMasc* mRNA in its sense orientation, included to show the sequence homologous to this mRNA present in each *Pxyfem* copy. CCCH ZFD = the N terminus zinc-finger domains characteristic of identified Masculiner homologues. CAEC = the ‘masculinizing’ domain characteristic of Masculinizer homologues. Lines cross over as *Pxyfem* copies are in the inverse orientation to the endogenous *PxyMasc* transcript. LINE = long interspersed element, LTR = long terminal repeat, TR= tandem repeat. Black box adjacent to exon 7 in Pxyfem long represents conserved 34bp sequence of unknown identification. Direction of *Pxyfem* transcription taken from observed RACE products. Direction of endogenous gene transcription taken from the ORF orientation of *PxyMasc* homologous sequence and surrounding identified ORFs. C: Phyologeny constructed using Clustal Omega including all 11 identified *Pxyfem* copies along with 250bp of up and downstream genomic flanking sequence.

The structure of the wider regions surrounding each *Pxyfem* locus varied dramatically. (Figure 5B). These areas contained a range of partial ORFs similar to transposable elements (TEs) as well as tandemly repeated elements which lacked discernible ORFs. Where present, these ORFs were orientated in the same direction as the *Pxyfem* loci direction of transcription (as deduced by RACE). Small RNA reads from the 12h embryo timepoint were found to map widely across the entire area of a searched c.22kbp genomic fragment which included *Pxfem* loci 1-4 (Supplementary Figure 1), though the density of antisense ssRNA reads mapping to this fragment increased dramatically in and around the four *Pxyfem* loci. This concurs with the situation in *B. mori* where the highly repetitive W-chromosome harbours large numbers of ‘transposable element graveyards’ where repeated and truncated copies of TE ORFs occur in a nested arrangement (6).

In contrast, the sequences immediately flanking the *Pxyfem* loci showed much higher conservation (Figure 5B). At the 3’ end of each *Pxyfem* (i.e. abutting truncated *PxyMasc* exon 4), a 251bp sequence was common to all 11 copies but absent elsewhere on this scaffold. These sequences contained an ORF which was aligned to generate an 82aa consensus sequence for BLASTp analysis. The consensus ORF showed closest similarity to the group-specific antigen (gag) polyproteins of LTR retrotransposons present in several lepidopteran families. The most similar hit (LOC105396011 with 51% aa identity over aligned region) and the highest number of significant hits (n=13) were found in the DBM male-derived assembly. The female-derived assembly was deliberately excluded from analysis to prevent the confounding effects of identifying *Pxyfem*-linked loci; hits in the male-derived assembly must be autosomal or Z-linked. Henceforth, we refer to *Pxyfem*-linked copies of this LTR retrotransposon as PLTR1, while the putative homologous endogenous gene we refer to as Endogenous LTR1 (ELTR1).

Similarly, immediately flanking the 5’ end of each short *Pxyfem* was a conserved c.256bp sequence (truncated to 108bp in one copy) whose translated ORF showed highest similarity (59.49% identity) to the gag polyprotein of a second LTR retrotransposon (LOC105389072) from the male-derived DBM genome (henceforth *Pxfem* linked LTR2 = PLTR2, putative endogenous homolog = ELTR2). Long *Pxyfem* also displayed an immediately adjacent conserved sequence but this was much shorter (34bp) and could not be identified. None of these conserved sequences were found elsewhere on this scaffold.

A phylogeny built using Clustal Omega (https://www.ebi.ac.uk/Tools/msa/clustalo/) which included each of the 11 *Pxyfem* copies as well as c. 250bp of up- and downstream genomic flanking sequence showed that these loci fell into two groups which could be differentiated by the presence of LTR2/absence of *PxyMasc* exon 7, or vice versa (Figure 5C). Interestingly, while *Pxyfem* copies within the same genomic cluster matched with each other in terms of their orientation, they did not necessarily group together within the phylogeny.

### *Pxyfem* and *fem* are functionally conserved

In *B. mori*, the *fem* piRNA targets *Masculinizer* transcripts for cleavage during embryogenesis (6). To assess whether *Pxyfem* plays a similar role in DBM we utilised a single transcript positive-readout system for identifying site-specific mRNA cleavage (31). This system uses two bacterial hairpins (CopT and CopA) placed in the 5’UTR and 3’UTR (respectively) of a luciferase expression construct, with an artificial polyA tail and downstream cleavage site placed upstream of CopA (Supplementary Figure 3A). In the absence of mRNA cleavage, the two hairpins function to partially suppress translation of the synthetic mRNA. If the target site is cleaved (e.g. by a ssRNA), the CopA hairpin is released from the transcript, exposing the synthetic polyA tail and allowing more efficient translation. Cleavage of a target mRNA sequence can thus be identified by changes in luciferase readout between treatments.

We adapted this design to assess whether the section of *PxyMasc* against which *Pxyfem* matched (exons 4-7) was sufficient to label a transcript for cleavage in DBM embryos, and whether the sex of those embryos influenced this behaviour. We observed a significant effect of embryo sex, with females showing a higher luciferase readout than males (p=0.0237, t=2.325, df=57, Figure 6). This result indicates that the presence of the W-chromosome (and thus the *Pxyfem* loci/expressed ssRNAs) is sufficient to cause the cleavage of transcripts bearing the *Pxyfem*-targeting *PxyMasc* sequence. This observation, paired with our previous result showing that these *PxyMasc* targeting ssRNAs are expressed in the early embryo at a time consistent with sex determination, strongly suggests the involvement of the *Pxyfem* system in determining female sexual fate in DBM through silencing *PxyMasc*.

**Figure 6:**
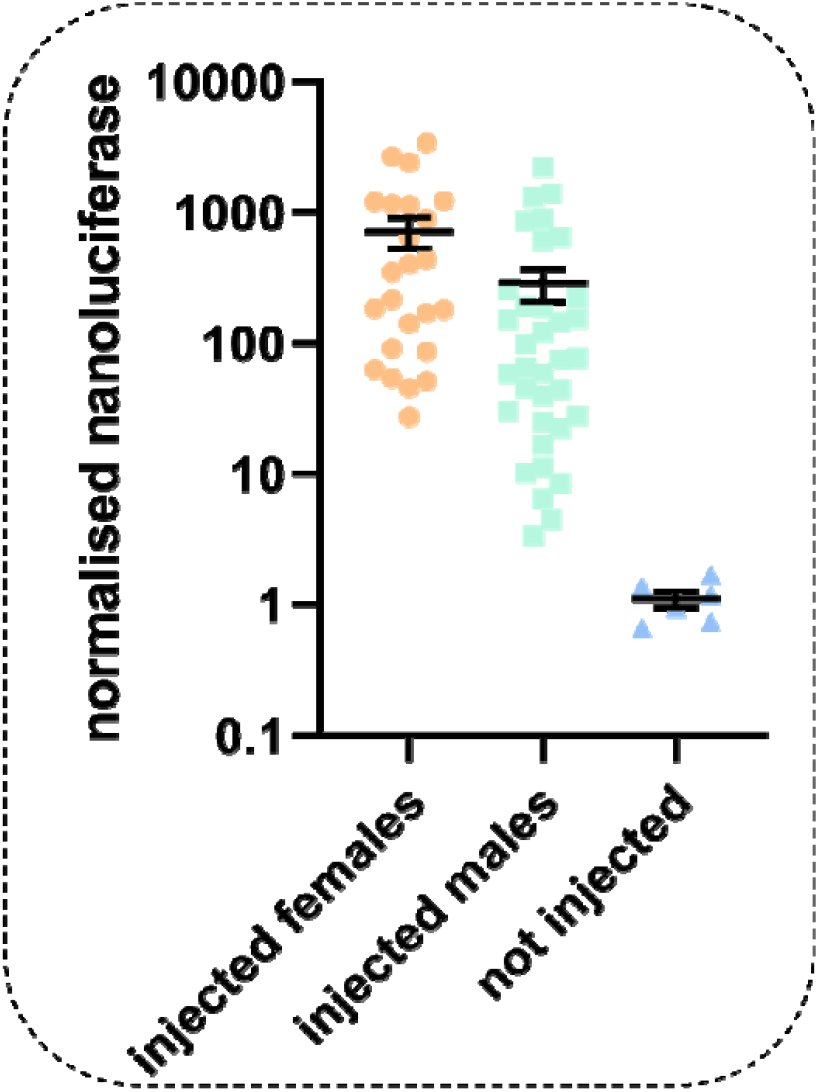
Female embryos display higher levels of cleavage induced positive readout signal than male embryos. X axis gives treatment, Y axis shows nanoluciferase readings in individual embryos (circles represent a single replicate embryo) normalised against firefly luciferase reading from the same embryo. Injected female embryos showed significantly higher levels of normalised nanoluciferase than injected males (P=0.0237, t=2.325, df=57, F= 24, M = 35, error bars = 95% CI), aligning with the hypothesis that cleavage of transcripts in this sex by *Pxyfem* derived ssRNAs increased subsequent translation.

### Origins of *Pxyfem*

The location of the *Pxyfem* loci on a large W-linked scaffold allows us to explore the possible origins of this system. Two interlinked questions include *How* and *When* such a mechanism might have arisen.

#### How did Pxyfem arise?

*Pxyfem* loci consist of partial *PxyMasc* cDNAs extending over several exon-exon boundaries (i.e. excluding three *PxyMasc* introns). This implies that the initial *Pxyfem* W-linkage event involved a spliced *PxyMasc* mRNA intermediate rather than a duplication of the Z-linked *PxyMasc* gene itself. In contrast, the 29bp *fem* in *B. mori* does not extend outside of *Masculinizer* exon 9 and thus no such conclusion can be drawn there.

Both *Pxyfem* and *fem* are in wider nested transposable and repeat element graveyards. LTR retrotransposons can drive the duplication of endogenous genes through a process known as retrotransposition (32, 33). These LTR retrotransposons increase their frequency through a ‘copy and paste’ mechanism in which their mRNA is reverse transcribed into cDNA, made double-stranded and then integrated elsewhere in the genome, forming a second copy of the original element (34). However, analysis in a diverse range of eukaryotes has found that mRNA template-switching events may occur during the reverse transcription step, allowing chimeric LTR retrotransposons to be generated (35). If the non-retrotransposon mRNA is derived from an endogenous gene, this may result in the partial duplication of that sequence, a ‘retrocopy’, if integration is successful. That, on the W-linked scaffold, the PLTR1 retrotransposon fragment was only found associated with *Pxyfem*, and its tight linkage to all 11 copies, suggests a role for an ancestral ELTR1 in the generation of *Pxyfem*. In *Drosophila melanogaster*, a hallmark of such template-switching retrotransposition events is short areas of microsimilarity at the ‘switch points’ between the LTR and endogenous gene transcripts (35). Consistent with this, at the precise putative switch point between ELTR1 and *PxyMasc* exon 4 we observed a shared sequence (ATTT) in these two transcripts whose length fits within the variance of such switch points observed previously in *D. melanogaster* retrocopies (Figure 7). We were unable to identify a sequence similar to ELTR1 on the opposite side of long *Pxyfem*, i.e. adjoining sequence homologous to *PxyMasc* exon 7, though a short conserved sequence of unknown origin was identified in this position (Figure 7A). Similarly, we were unable to identify the LTR sequences that would have been required for genomic integration of a chimeric retrotransposon. Sequence drift in these areas, as well as rounds of recombination and reduplication may have removed these sequences or altered them beyond recognition. Previous studies exploring gene duplication through retrotransposition have utilised ‘young’ retrocopies specifically to minimise these confounding effects (35). The severely truncated nature of the TE ORFs identified on this W-chromosome scaffold implies that this degeneration and fragmentation would not have been unique to the hypothesised ancestral *Pxyfem*-PLTR1 integration.

**Figure 7:**
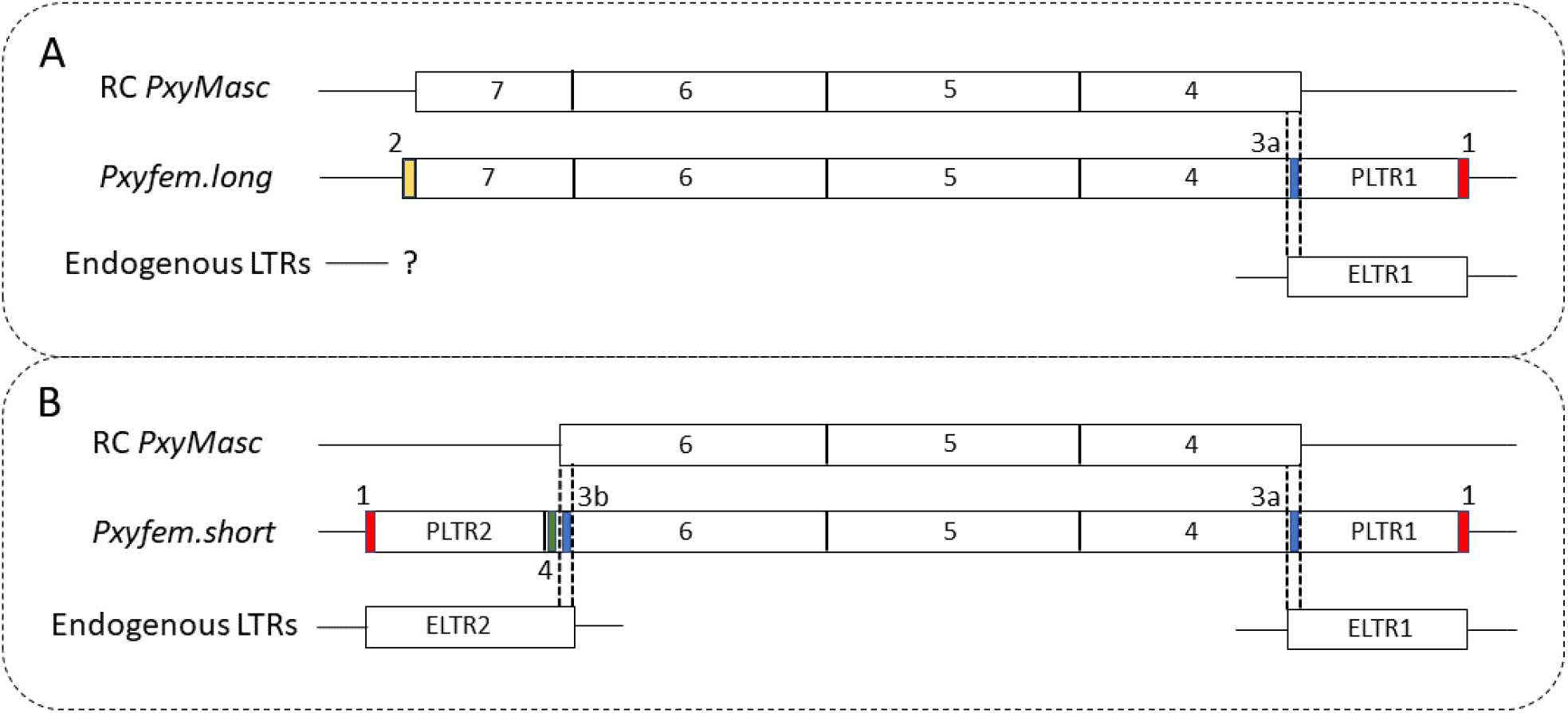
Microsimilarity and repeat regions suggest a possible role for ‘retrotransposition’ in the origins of *Pxyfem*. A: Analysis of ‘*Pxyfem* long’. From top to bottom, the three sequence are - RC *PxyMasc* = Reverse complement of *PxyMasc* transcript. *Pxyfem.long* = generalised schematic of the *Pxyfem.long* copies, including 250bp of up and downstream genomic flanking sequence. Endogenous LTRs = the closest homologue to PLTR1 which flanks *Pxyfem.long*. Numbers above coloured segments in *Pxyfem.long* denote conserved sequences as follows. 1 =10bp repeat : (ACACAAGGCT), **2** =34bp conserved but unidentified sequence : (ACCCCAGCAGTTACATGTTGGGAAGCAGAAATTT). **3a** = *PxyMasc*/<u>ELTR1</u> switch point : (ATTT). Dashed lines represent the putative switch points between RC *PxyMasc* and ELTR1 which gave *Pxyfem*. B: Analysis of ‘*Pxyfem* short’. Three sequences same as in A but exchanging the generalised *Pxyfem.long* sequence with the generalised *Pxyfem.short* sequence. **3b** = *PxyMasc*/<u>ELTR2</u> switch point : *PxyMasc* sequence = AA<u>GA</u>AG, ELTR1 sequence = AA<u>CC</u>AG, putative switch point = AA<u>GC</u>AG). Dashed lines represent the putative switch points between RC *PxyMasc* and ELTR2 which gave *Pxyfem.short*. **4** = 15bp tandem duplication : (<u>AGTCTGCTGGTTAAG</u>AAGA<u>AGTCTGCTGGTTAAG</u>).

The architecture of short *Pxyfem* provides a clue as to how these reduplication events may have occurred. Relative to long *Pxyfem*, the four short *Pxyfem* copies are truncated on their 5’ ends and in place of this lost sequence contain the PLTR2 retrotransposon ORF fragment (Figure 7B). The putative switch point between short *Pxyfem* and PLTR2 is more difficult to assess than for PLTR1 due to a 15bp tandem duplication of the terminal *Pxyfem* sequence at this junction. However, an area of potential microsimilarity was identified in the *PxyMasc* CDS immediately downstream of this duplicated sequence and immediately upstream of the PLTR2 homologous region in ELTR2, with the putative switch point containing a combination of the SNPs in these two regions). Short *Pxyfem* may represent the result of a retrotransposition event involving an ancestor of the ELTR2 retrotransposon and the previously integrated PLTR1-long *Pxyfem* fusion. Consistent with the hypothesised more recent origin of short *Pxyfem*, we observed a 10bp element repeated at the junctions of PLTR1/PLTR2 and their respective flanking genomic regions which may represent the target site duplication (TSD) signal of a LTR retrotransposition event (34).

#### When did Pxyfem arise?

There are two mutually exclusive scenarios regarding the evolutionary relationship between the *B. mori* and *P. xylostella ‘feminizer’* systems. Scenario 1: *fem* and *Pxyfem* derive from a common ancestor (‘proto-*feminizer*’) that evolved prior to the split of the *Plutellidae* from the rest of the Ditrysisa, c.125MYA. Scenario 2: *fem* and *Pxyfem* evolved independently after this split and the functional similarities between them represent convergent evolution. These two scenarios are explored below.

Assuming that *Pxyfem* is a partial *PxyMasc* retrocopy, one way to assess the two scenarios is through comparison of the ‘fossilised’ TEs involved – PLTR1 and PLTR2 - with other lepidopteran LTR retrotransposons. Under scenario 1 (common origin) we would *not* expect these to show higher similarity to remaining ancestral sequences in the DBM genome (i.e. ELTR1/ELTR2) than to homologues of those sequences in other higher lepidopterans. Conversely, if the original template switching event took place *after* the radiation of the Ditrysians (scenario 2 – independent origin) we *would* expect that PLTR1 and PLTR2 would resemble ELTR1/ELTR2 more than other lepidopteran homologues. Consistent with scenario 2, when BLASTed, translated PLTR1/PLTR2 consensus sequences show higher similarity to LTR retrotransposons in the male-derived DBM genome (i.e. in DBM autosomal or Z-linked regions) than to hits in other lepidopteran assemblies. Moreover, we could not identify sequences similar to any retrotransposable element ORFs in the 767bp *B. mori fem*-precursor transcript. Though these LTR-related sequences (i.e a hypothetical BLTR1/BLTR2) may have been lost from the *B. mori fem* transcript through sequence turnover, it does not explain why the remnants of these sequences in DBM (PLTR1/PLTR2) contain ORFs more similar to those found in DBM autosomal regions.

In principle, this greater similarity could be due to homogenisation of paralogous sequences by non-allelic gene conversion (NAGC). However, the nucleotide sequences of the PLTR1 copies and ELTR1 (i.e. the sequences on which the homology-based NAGC would have theoretically acted) show relatively low similarity (PLTR1/ELTR1 = c. 57% identity over c. 251bp, PLTR2/ELTR2 = c. 60% identity over 236bp). It would appear that gene conversion, which is known to convert runs of on average a few hundred base pairs at a time, at a rate 10-100x faster than point mutation (36), has had little observable role in the concerted evolution of these PLTR/ELTR pairs.

The two potential origin scenarios should also have different implications for evolution of *PxyMasc* and *Pxyfem* sequences. The function of *Pxyfem* established here is to provide a source of ssRNAs targeting *PxyMasc*. We would expect, therefore, that selection pressure on differing regions of *Pxyfem* to maintain nucleotide agreement with *PxyMasc* would be in proportion to the relative contributions of those areas towards that function. We would further anticipate that the redundancy of the 11 *Pxyfem* copies may have allowed the sequences of ‘low-functionality’ areas to drift considerably over evolutionary time, both within and between copies. Under the ancient origin of scenario 1, its high level of within-sequence agreement would thus suggest that there has been extremely strong selection pressure across the *Pxyfem* sequence to maintain this agreement. This expectation is difficult to match with the observation that the area of *Pxyfem* which our sequencing identified as producing ssRNAs against *PxyMasc* is relatively concentrated. Indeed, the first 46bp of *Pxyfem* short and *Pxyfem* long, despite showing 89% nucleotide identity to exon 4 of *PxyMasc*, did not produce a single antisense read mapping to *PxyMasc*. Similarly, for *Pxyfem* long, the final 97bp, which displayed 96% nucleotide identity with exons 6 and 7 of *PxyMasc*, produced only a single antisense read mapping to *PxyMasc*. Given that these two regions appear to contribute few or no ssRNAs to the silencing of *PxyMasc*, it is difficult to explain why such tight sequence agreement between them and the relevant regions of *PxyMasc* would have been maintained for > 125 million years.

Under scenario 2 (independent origin) this situation can be explained easily as here the sequence-specific *feminizer* systems of DBM and *B. mori* arose *after* the sequence divergence of their two respective *Masculinizer* homologues. Under this scenario, both the observed within-system agreement and the retention of ‘low-functionality’ areas of *Pxyfem* are a consequence of the relatively recent origins of this system. This seems to fit the available data better than scenario 1.

In conclusion, we have identified a W-linked multi-copy ssRNA generating system in DBM which targets the male determining gene *PxyMasc*. Consistent with the hypothesis that this system functions like *feminizer* in *B. mori* to silence the *Masculiniser* gene and determine female sexual fate, we observed that it initiates expression during early embryogenesis, peaking 12h post laying. Previously, we established that the *PxyMasc* transcript - initially expressed (3-6h post laying) in both male and female embryos - can no longer be observed in female embryos by 24h post laying, correlating with a transition to female-specific splicing of doublesex. The peak expression of the *Pxyfem* system thus overlaps with the reduction in *PxyMasc* transcripts observed in female DBM embryos and a transition towards female somatic differentiation (the function of female-form doublesex). Further supporting the proposed mode of action of *Pxyfem*, experiments with exogenous transcripts ‘labelled’ with the *Pxyfem* target sequence indicated that this 325bp sequence is targeted for ssRNA-mediated cleavage in the early female embryo. Intriguingly, our analyses suggest the more likely origin scenario for this *Pxyfem* system is that it arose after the divergence of the basal *Plutellidae* from the higher Ditrysia, and that similarities between it and *B. mori fem* represent convergent evolution. Our model for the origin and evolution of *Pxyfem* implicates involvement of LTR-transposon retrotransposition. To our knowledge this is the first recorded example of such an event having generated a component of a sex determination system, let alone one which may represent a master regulator of sexual fate. Further work elucidating the evolutionary relationship between the lepidopteran W-chromosomes and, especially, the sequencing of other members of the *Plutellidae*, may further aid in this analysis.

## Methods

### Insect details

DBM (originally collected from Vero Beach, Florida, USA) were reared on beet armyworm artificial diet (Frontier Biosciences, Germantown, Maryland, USA) under a 16:8⍰h light : dark cycle, 25⍰°C and 50% relative humidity.

### Primers

Details of all primers used in this study can be found in Supplementary table 2.

### Generating DBM small RNA libraries and mapping to *PxyMasc*

Larval L1 samples: Newly hatched (within 1 h) DBM larvae were individually collected, placed in RNA*later* (Thermo Fisher Scientific, Waltham, Massachusetts, USA), immediately frozen in liquid nitrogen and stored at -80⍰°C. RNA was extracted from each sample using a miRNeasy Mini Kit and a RNeasy MinElute Cleanup Kit (both from (Qiagen, Valencia, CA, USA)), in order to isolate both >200nt and <200nt fractions from each sample. <200nt isolates were immediately frozen at -80⍰°C while >200nt isolates were used as template for cDNA synthesis (LunaScript RT Mastermix Kit, New England Biolabs, MA, USA) and subsequent *doublesex* sexing PCR as previously described (25). Once the sex of each sample had been confirmed through observing *doublesex* splicing patterns, <200nt samples were combined to make a male (n=7) and female (n=5) small RNA isolate. These two combined samples were ethanol precipitated and used to generate male and female L1 small RNA libraries using the NEBNext Multiplex Small RNA Library Prep Set for Illumina (New England Biolabs, MA, USA). Indexed libraries were size selected for sRNAs (<150nt) using AMPure XP magnetic beads (1.3X followed by 3.7X) (Beckman Coulter, Brea, California, USA). The final size of each library was determined using a D100 High Sensitivity Screentape on a TapeStation 2200 (Agilent, Santa Clara, California, USA). Libraries were quantified using the Qubit dsDNA BR assay (ThermoFisher, MA, USA), diluted to 20nM and further quantified using the NEBNext Library Quant Kit for Illumina (New England Biolabs, MA, USA). Samples were pooled and diluted to 4nM and run on an Illumina Miseq by the Bioinformatics, Sequencing and Proteomics Facility at The Pirbright Institute.

#### Embryo samples

Adult DBM were allowed to lay eggs on a cabbage juice painted parafilm sheet for 5 minutes. Laid eggs were removed from the cage after this time and placed into a humidified petri dish. At each time point (3, 6, 9, 12 and 24h post laying) 100 eggs were collected from the parafilm sheet, pooled, placed in RNA*later* and frozen in liquid nitrogen. Downstream processing of these 5 egg samples followed the same workflow outlined above, however, >200nt isolates were collected but not used for *doublesex* sexing through RT-PCR as these were mixed-sex collections/extractions. Library preparation and analysis were performed as for the L1 samples. The prepared small RNA libraries were sequenced using the Illumina Miseq platform as for the L1 samples.

#### Sequencing analysis

Sequenced reads were checked for quality using FastQC (37). The TruSeq sequencing adapters (RNA 5’ adapter: GUUCAGAGUUCUACAGUCCGACGAUC; (RA5); part # 15013205, RNA 3’ adapter: TGGAATTCTCGGGTGCCAAGG; (RA3); part # 15013207) and the poor-quality reads (Phred scores; Q<20) were removed using Trimmomatic (38). Further, the reads which were not in the ssRNA size range (20-40bp) were filtered out using the Seqkit toolkit (39). The pre-processed reads were then mapped to the *PxyMasc* sequence using GSNAP (40). The mapped alignment file was processed to identify sense and antisense reads using SAMtools (41). Mapped outcomes (size distribution and density plots) from different time-course experiments and male-female-specific libraries were then plotted using the Graphpad Prism 8 (GraphPad Software Inc., USA). Integrated Genomics Viewer (IGV) (42) was used to visualise the alignment files. The nucleotide usage (consensus sequences) from the identified similar size ssRNAs were plotted as sequence logos using Weblogo3 (43).

### PxyFem RACE

From small RNA mapping of reads deriving from L1 samples, *PxyMasc* appeared to be targeted by two ssRNA groups being complementary to exons 4 and 5/6, respectively. These two areas were used to design 5’ and 3’ RACE primers (see Supplementary Table 2) in order to identify mRNA sequence from the *Pxyfem* precursor transcript. 5’ and 3’ RACE ready cDNA was generated from a pooled female pupal RNA sample and also from a pooled adult female ovary sample (RNA extracted using RNeasy MinElute kit, RACE conducted using SMARTer 5⍰/3⍰ RACE kit - Takara Bio, Kyoto, Japan). Visible bands were cloned using the NEB PCR cloning kit (New England Biolabs, MA, USA) and sanger sequenced. Sequences which did not show evidence of mispriming (i.e. those that did show significant homology to *PxyMasc* in reverse complement) were further characterised through BLASTn and translated reading frames through BLASTp analysis (https://blast.ncbi.nlm.nih.gov/Blast.cgi). For BLASTn Whole-genome Shotgun Contigs (WGS) of organism ‘*Plutella xylostella*’ were used as search subject. For BLASTp, non-redundant protein sequences (nr) of the same organism was used. In the latter case, significant protein matches were subsequently re-blasted without species limitations in order to identify the families to which these belonged. Putative domains within protein homologs were identified using the NCBI conserved domain search (https://www.ncbi.nlm.nih.gov/Structure/cdd/wrpsb.cgi).

### Assesment of *Pxyfem*-W chromosome linkage

gDNA from 6 male and 5 female pupae (sexed by eye) was extracted using the NucleoSpin Tissue kit (Macherey-Nagel, Düren, Germany), pooled by sex and used as template for subsequent PCRs. Four PCRs were run on each template, informed by the sequencing results arising from 5’ RACE analysis. PCR set 1 (primers LA4886 + LA4887) was designed to amplify across exons 5 and 6 of the region showing sequence homology to *PxyMasc*, PCR set 2 (primers LA4888 + LA4889) was designed to amplify from the internal region (IR) of the transcript across the region showing sequence homology to exons 6 and 5 of *PxyMasc* (and thus has the same reverse primer as primer set 1), PCR set 3 (primers LA4890 + LA4891) was designed within the Multiply Repeated Region 1 (MRR1), PCR set 4 (primers LA309 + LA310) was designed to amplify within the 17S genomic locus as a positive gDNA control). All PCRs were run using DreamTaq DNA polymerase (Thermo Fisher Scientific, MA, USA) and the programme 98⍰°C – 1 min, 35⍰cycles of 98⍰°C – 30⍰s, 50⍰°C – 30⍰s, 72⍰°C – 30⍰s, final extension 72⍰°C – 21min. A no template control (NTC) was also run for each primer set. Primers sequences are detailed in Supplementary table 2.

### Positive readout mRNA cleavage experiment

The positive readout cassette was based on a previously demonstrated design and was synthesised by Genewiz Inc. In brief, the 325bp *PxyMasc* sequence matching to *Pxyfem* was placed directly downstream of a polyA sequence and directly upstream of the CopA sequence. As previously, CopT was placed in the 5’UTR of nanoluciferase. This nanoluciferase transcript was placed under the control of the Hr5/Ie1 promoter and this cassette was cloned into a pre-existing backbone (AGG1938) which contained an Op/Ie2-ZsGreen-Sv40 fluorescent marker cassette to give the positive readout construct AGG2208 (Supplementary Figure 3A – Genbank accession number (pending)). This plasmid was injected into <30min old DBM embryos alongside a second plasmid expressing firefly luciferase also under the control of Hr5/Ie1 (AGG1186 - Supplementary Figure 3B - Genbank accession no. MT119956 (44)), to act as a normalisation control. After 48 hours, individual embryos were sexed by genomic (W-chromosome) PCR and normalised nanoluciferase expression calculated.

Prior to the experiment, AGG2208 and AGG1186 were combined with molecular grade H_2_0 (final plasmid concentrations of 400ng/ul and 300ng/ul, respectively) to provide an injection mix. This mix was injected into <30min old DBM embryos using techniques previously described (45). 48h later, embryos were inspected under a fluorescence microscope and eggs which did not show strong evidence of ZsGreen fluorescence (the fluorophore central marker on the injected plasmid which will fluoresce independently of any other effects) were excluded on the basis that these either had little to no mix injected, or had died at an early stage after injection. Remaining eggs were transferred into individual PCR tubes and frozen at -20 °C.

Subsequently, eggs were individually homogenised in 5ul DEPC water within the PCR tube and 1ul was used for gDNA extraction using Phire Animal Tissue Direct PCR Kit (ThermoFisher, MA, USA) according to manufacturer’s instructions, following the dilution protocol but adding 1ul of the homogenised sample instead of tissue. Two different PCRs were performed for each egg sample, one for a confirmed autosomal locus (Multiply Repeated Region 1 - MRR1) to act as an extraction control (primers LA4890 + LA4891), and one for a confirmed W-specific locus (*Pxyfem*) to identify female embryos (primers LA4888 + LA4889). These reactions were confirmed in the ‘Assessment of *Pxyfem* W-chromosome linkage’ section, above. PCRs were performed using Phire Animal Tissue Direct PCR Kit according to manufacturer’s instructions using 2ul of gDNA template for the genomic PCR and 3ul of template for the female specific PCR. PCRs were performed using a standard cycling heat block with the following cycler program: 98⍰°C – 5 min, 35⍰cycles of 98⍰°C – 5⍰s, 64.9 (MRR1) / 65.4 (*Pxyfem*)⍰°C – 5⍰s, 72⍰°C – 20 s, final extension 72⍰°C – 1⍰min. Primer sequences are detailed in Supplementary table 2.

To the remaining 4ul of each egg homogenate sample, passive lysis buffer (Promega, WI, USA) was added to a 1x final dilution and incubated for 15min at room temperature. The resulting product was used to perform luciferase assays using Nano-Glo® Dual Reporter Assay kit (Promega, WI, USA) according to manufacturer’s instructions. Absorbance was measured using a GloMax Microplate reader (Promega, WI, USA). Both firefly and nanoluciferase values were measured and nanoluciferase values of each sample were normalised against corresponding firefly values. Analysis was conducted on Prism (Graphpad, San Diego, California, USA) using a two-tailed, unpaired t-test. Female embryo replicates=24, male embryo replicates=35.

### Assessment of *Pxyfem* operon linkage

Samples included 1) cDNA generated from >200nt RNA extracted previously from the 12h embryo sample (LunaScript RT Mastermix Kit, New England Biolabs). 2) A no-RT control of the embryo sample. 3) Molecular grade H20. 4) Female pooled gDNA extracted previously for the ‘Assessment of *Pxyfem* W chromosome linkage’ experiment. Each of these samples was used as template for a PCR using primers LA2549 and LA4887 (Supplementary table 2) which extended ‘outwards’ from each *Pxyfem* locus. As such, amplicons produced by these reactions must occur between *Pxyfem* loci. PCRs were performed using Q5 polymerase (New England Biolabs, MA, USA) and the following thermocycler programme 98⍰°C – 1 min, 35⍰cycles of 98⍰°C – 30⍰s, 68⍰°C – 30⍰s, 72⍰°C – 3 min, final extension 72⍰°C – 2⍰min.

## Acknowledgements

DKP’s PhD studentship was funded by The Pirbright Institute. LC was supported by a Wellcome Trust grant made to LA [110117/Z/15/Z]. For the purpose of Open Access, the author has applied a CC BY public copyright license to any Author Accepted Manuscript (AAM) version arising from this submission. MAEA and LA were funded through a Defense Advanced Research Projects Agency (DARPA) award [N66001-17-2-4054] to Kevin Esvelt at MIT. The views, opinions and/or findings expressed are those of the authors and should not be interpreted as representing the official views or policies of the U.S. Government. VCN and THS were supported by European Union H2020 Grant nEUROSTRESSPEP (634361) made to LA. XX was supported by the CSC Scholarship from the Chinese Government and a PhD student exchange program from Fujian Agriculture and Forestry University (FAFU). THS was supported by a UK Biotechnology and Biological Sciences Research Council (BBSRC) Impact Acceleration Account grant (BB/S506680/1) made to THS and LA. LA and THS were additionally supported by core funding from the BBSRC to The Pirbright Institute (BBS/E/I/00007033, BBS/E/I/00007038 and BBS/E/I/00007039). CR was supported by a Wellcome Trust grant made to LA (200171/Z/15/Z). The authors would like to acknowledge the assistance of Dr Graham Freimanis at the Pirbright Institute Bioinformatics, Sequencing and Proteomics Unit.

## Notes

### Competing Interest Statement

The authors have declared no competing interest.

