## supporting info for "Small silencing RNAs expressed from W-linked retrocopies of *Masculinizer* target the male-determining gene *PxyMasc* during female sex determination in the Diamondback moth *Plutella xylostella*"

**Supporting Information**

Table 1: **Summary of small RNA deep sequencing and mapping to *PxyMasc* mRNA sequence**

| **Library** | **No. of reads post QC / Trimming (<= 40bp length)** | **No. of mapped reads** | **Mapped forward reads** | **Mapped reverse reads** | **Mapped reads/million total reads** |
| --- | --- | --- | --- | --- | --- |
| **Female L1** | **3328300** | **36** | **12** | **26** | **10.81633266** |
| **Male L1** | **2369543** | **2** | **2** | **0** | **0.844044611** |
| **3h embryos** | **4814110** | **0** | **0** | **0** | **0** |
| **6h embryos** | **4617092** | **36** | **26** | **10** | **7.797115587** |
| **9h embryos** | **4284613** | **146** | **88** | **58** | **34.07542291** |
| **12h embryos** | **4090068** | **180** | **91** | **89** | **44.00904826** |
| **24h embryos** | **3741120** | **124** | **51** | **73** | **33.14515439** |

Table 2: **Primers used in this study.**

| **Primer** | **sequence** |
| --- | --- |
| **LA2549** | GCTCCCTTCGCGGCCTTAC |
| **LA4890** | CGAGTATGAGGAATTAAACAGCCTCC |
| **LA4891** | GAAGATATTCCGGACGTCAGTCTAG |
| **LA4888** | CATTAGTGGTATCTAATAGGGATCTCAACTG |
| **LA4889** | CGCAGTCTGCTGGTTAAGAAGAAG |
| **LA4886** | CTTCTTCTTAACCAGCAGACTGCG |
| **LA4887** | CAGCTCAAGACAAGAGTTGTTGAGAG |
| **LA309** | CAAGCCTCTTCGTAACAAGATCG |
| **LA310** | CAGCTTGATGGAGATACCACGC |
| **5’ RACE primer R1** | GCCACGCCGAGTAAGGCCGCGAAGGG |
| **5’ RACE primer R2** | CTGAATGTGCTCAGCTCAAGACAAGA |
| **3’ RACE primer R1** | CCCTTCGCGGCCTTACTCGGCGTGGC |
| **3’ RACE primer R2** | TCTCAACAACTCTTGTCTTGAGCTGAGC |

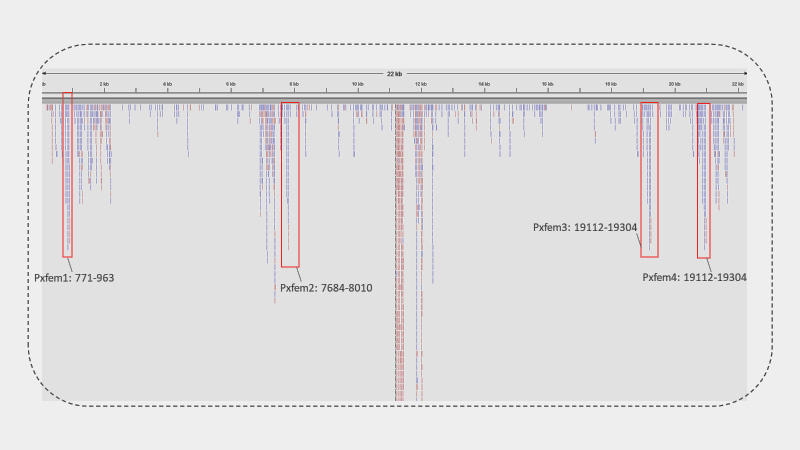

Figure 1: **Pxyfem loci occur within clusters of high ssRNA expression.** Graphic downloaded from Integrative Genomics Viewer showing reads from female L1 pool mapped to a 22Kb fragment of the CABWKK010000004 genomic scaffold encompassing the first *pxyfem* cluster (*pxyfem* 1-4). Approximate locations of the *pxyfem* loci shown. Blue read colour signifies reads mapping to reverse strand. Red read colour signifies mapping to forward strand. The four *pxyfem* loci occur with in areas with high levels of reverse strand mapping reads (i.e. producinig antisense ssRNAs against *PxyMasc* mRNA).

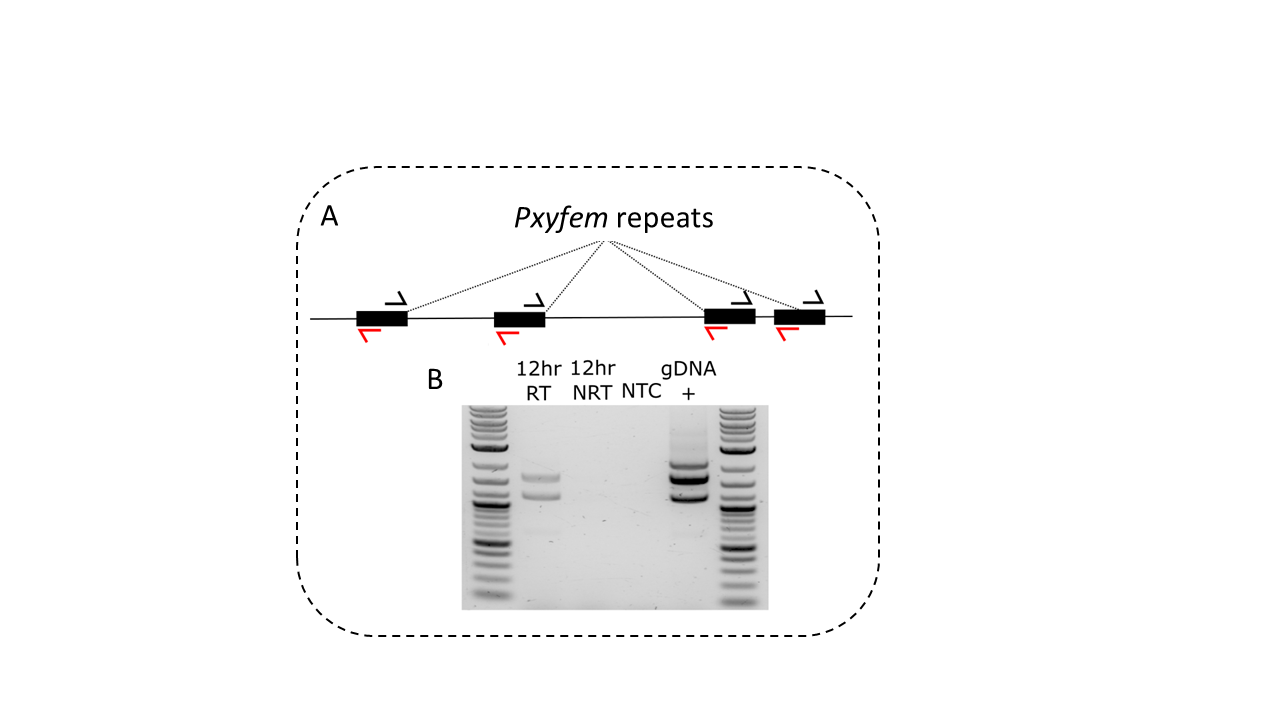

Figure 2: **Closely-linked *Pxyfem* copies are expressed on the same transcript.** A: Schematic showing four hypothetically placed *Pxyfem* copies. Primer symbols represent forward and reverse primers used in B which were conserved amongst *Pxyfem* copies. B: Results of PCR using primers shown in A. From left to right - 12h RT = RT-PCR using cDNA from 12h embryos. 12h NRT = No reverse transcriptase control of first lane. NTC = no template control for PCR in lane 4. gDNA+ = PCR using pooled female DBM gDNA. For the RT-PCR, bands would only be expected if different *Pxyfem* copies occurred on the same transcript. The presence of two bands suggests at least three *Pxyfem* copies are expressed on the same transcript.

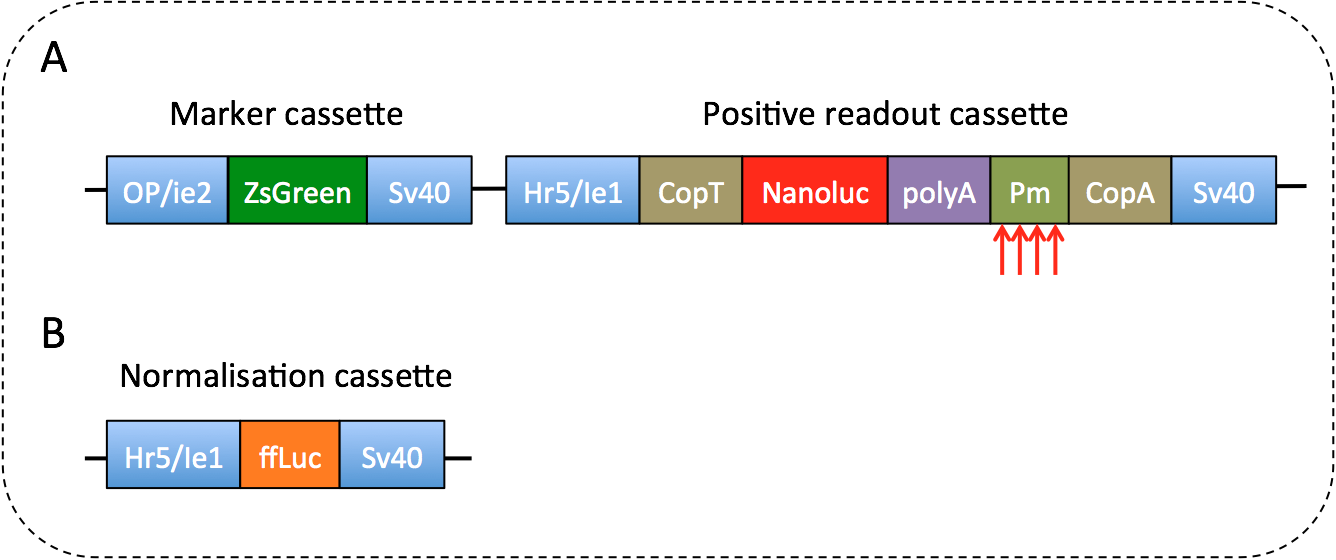

Figure 3: **Constructs used for positive readout cleavage experiment.** A) Schematic of AGG2208. In the absence of transcript cleavage, the bacterial hairpins CopT and CopA dampen translation of the positive readout mRNA transcript. In the presence of transcript cleavage (signified by the red arrows) the CopA inhibitory hairpin is removed and the internal, synthetic polyA tail exposed, allowing increased translation of the transcript and higher nanoluciferase signal to be observed. ‘Pm’ represents the 325bp sequence of *PxyMasc* against which *Pxyfem* matches (i.e. the putative *PxyMasc* target site of *Pxyfem*). B) Schematic of AGG1186. ‘ffLuc’ represents the firefly luciferae ORF.
